# Clustering trees: a visualisation for evaluating clusterings at multiple resolutions

**DOI:** 10.1101/274035

**Authors:** Luke Zappia, Alicia Oshlack

## Abstract

Clustering techniques are widely used in the analysis of large data sets to group together samples with similar properties. For example, clustering is often used in the field of single-cell RNA-sequencing in order to identify different cell types present in a tissue sample. There are many algorithms for performing clustering and the results can vary substantially. In particular, the number of groups present in a data set is often unknown and the number of clusters identified by an algorithm can change based on the parameters used. To explore and examine the impact of varying clustering resolution we present clustering trees. This visualisation shows the relationships between clusters at multiple resolutions allowing researchers to see how samples move as the number of clusters increases. In addition, meta-information can be overlaid on the tree to inform the choice of resolution and guide in identification of clusters. We illustrate the features of clustering trees using a series of simulations as well as two real examples, the classical iris dataset and a complex single-cell RNA-sequencing dataset. Clustering trees can be produced using the clustree R package available from CRAN (https://CRAN.R-project.org/package=clustree) and developed on GitHub (https://github.com/lazappi/clustree).

## Introduction

Clustering analysis is commonly used to group similar samples across a diverse range of applications. Typically, the goal of clustering is to form groups of samples that are more similar to each other than to samples in other groups. While fuzzy or soft clustering approaches assign each sample to every cluster with some probability, and hierarchical clustering forms a tree of samples, most methods form hard clusters where each sample is assigned to a single group. This goal can be achieved in a variety of ways, such as by considering the distances between samples (e.g. *k-*means [1-3], PAM [4]), areas of density across the dataset (e.g. DBSCAN [5]) or relationships to statistical distributions [6].

In many cases the number of groups that should be present in a dataset is not known in advance and deciding the correct number of clusters to use is a significant challenge. For some algorithms, such as *k-*means clustering, the number of clusters must be explicitly provided. Other methods have parameters that, directly or indirectly, control the clustering resolution and therefore the number of clusters produced. While there are methods and statistics (such as the elbow method [7] or silhouette plots [8]) designed to help analysts decide which clustering resolution to use, they typically produce a single score which only considers a single set of samples or clusters at a time.

An alternative approach would be to consider clusterings at multiple resolutions and examine how samples change groupings as the number of clusters increases. This has lead to a range of cluster stability measures [9], many of which rely on clustering of perturbed or sub-sampled datasets. For example, the model explorer algorithm sub-samples a dataset multiple times, clusters each sub-sampled dataset at various resolutions and then calculates a similarity between clusterings at the same resolution to give a distribution of similarities which can inform the choice of resolution [10]. One cluster stability measure that isn’t based on perturbations is that contained in the SC3 package for clustering single-cell RNA-sequencing data [11]. Starting with a set of cluster labels at different resolutions each cluster is scored, with clusters awarded increased stability if they share the same samples as a cluster at another resolution, but penalised for being at a higher resolution.

A similar simple approach is taken by the clustering tree visualisation we present here, without calculating scores: (i) a dataset is clustered using any hard clustering algorithm at multiple resolutions, producing sets of cluster nodes, (ii) the overlap between clusters at adjacent resolutions is used to build edges, (iii) the resulting graph is presented as a tree. This tree can be used to examine how clusters are related to each other, which clusters are distinct and which are unstable. In the following sections we describe how we construct such a tree and present examples of trees built from a classical clustering dataset and a complex single-cell RNA-sequencing (scRNA-seq) dataset. The figures shown here can be produced in R using our publicly available clustree package. Although clustering trees can not directly provide a clustering resolution to use they can be a useful tool for exploring and visualising the range of possible choices.

## Building a clustering tree

To build a clustering tree, we start with a set of clusterings allocating samples to groups at several different resolutions. These could be produced using any hard-clustering algorithm that allows control of the number of clusters in some way. For example, this could be a set of samples clustered using *k-*means with *k* =1, 2, 3 as shown in Figure 1. We sort these clusterings so that they are ordered by increasing resolution (*k*), then consider pairs of adjacent clusterings. Each cluster *c_k,i_* (where *i* = 1, *…,n* and *n* is the number of clusters at resolution *k*) is compared with each cluster *c_k+i,j_* (where *j =* 1*,…, m* and *m* is the number of clusters at resolution *k* +1). The overlap between the two clusters is computed as the number of samples that are assigned to both *c_k,i_* and *c_k+i,j_*. We next build a graph where each node is a cluster and each edge is an overlap between two clusters. While we refer to this graph as a tree in this paper for simplicity it can more correctly be described as a polytree, a special case of a directed acyclic graph where the underlying undirected graph is a tree [12].

**Figure 1:**
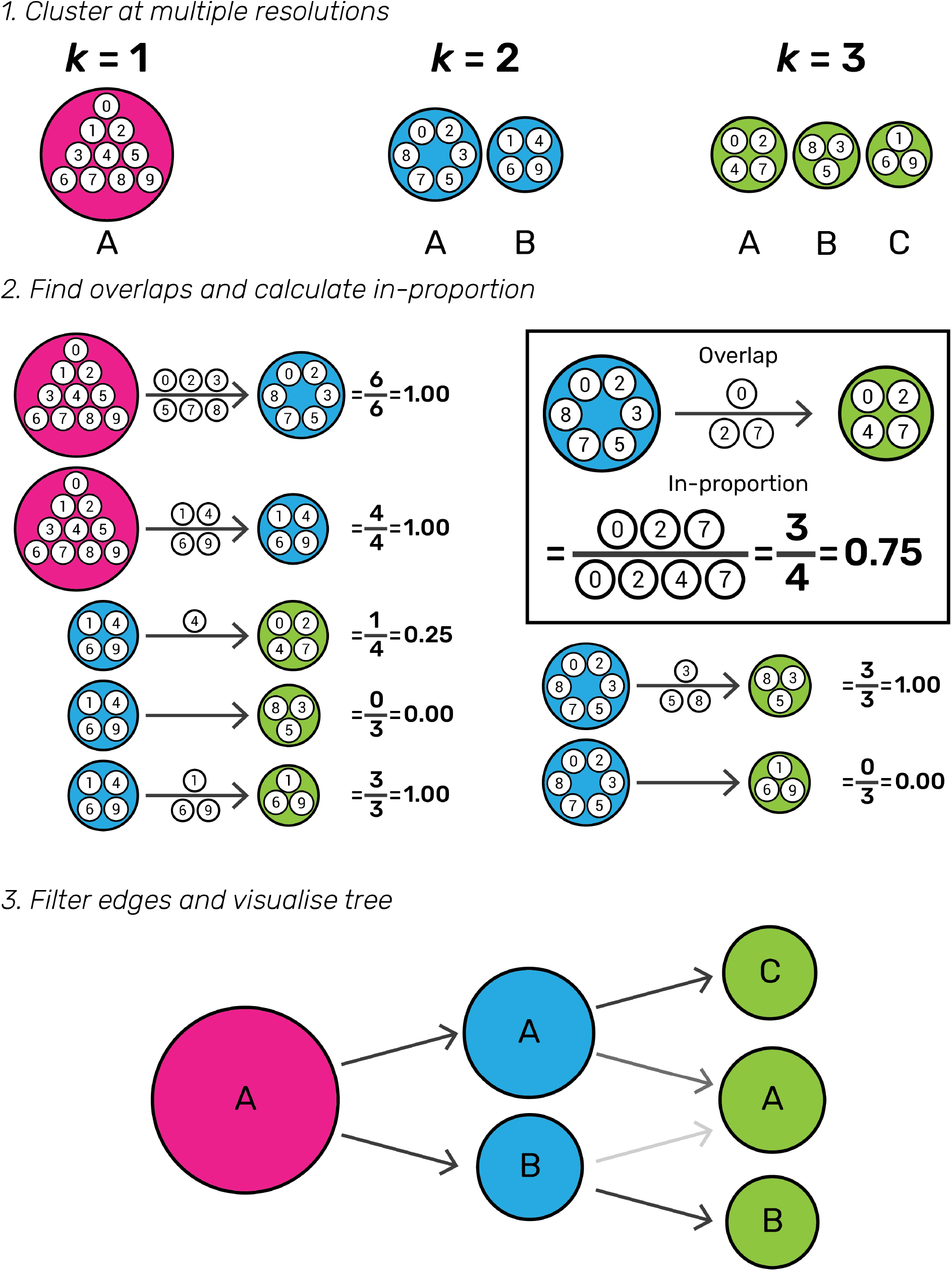
Illustration of the steps required to build a clustering tree. First a dataset must be clustered at different resolutions. The overlap in samples between clusters at adjacent resolutions is computed and used to calculate the in-proportion for each edge. Finally the edges are filtered and the graph visualised as a tree.

Many of the edges will be empty, for example in Figure 1 no samples in Cluster A at *k =* 2 end up in Cluster B at *k* = 3. In some datasets there may also be edges that contain few samples. These edges are not informative and result in a cluttered tree. An obvious solution for removing uninformative, low-count edges is to filter them using a threshold on the number of samples they represent. However, in this case the count of samples is not the correct statistic to use because it favours edges at lower resolutions and those connecting larger clusters. Instead we define the in-proportion metric as the ratio between the number of samples on the edge and the number of samples in the cluster it goes towards. This metric shows the importance of the edge to the higher resolution cluster independently of the cluster size. We can then apply a threshold to the in-proportion in order to remove less informative edges.

The final graph can then be visualised. In theory any graph layout algorithm could be used but for the clustree package we have made use of the two algorithms specifically designed for tree structures available in the igraph package [13]. These are the Reingold-Tilford tree layout, which places parent nodes above their children [14], and the Sugiyama layout which places nodes of a directed acyclic graph in layers while minimising the number of crossing edges [15]. Both of these algorithms can produce attractive layouts and as such we have not found the need to design a specific layout algorithm for clustering trees. By default the clustree package uses only a subset of edges when constructing a layout, specifically the highest in-proportion edges for each node. We have found that this often leads to more interpretable visualisations, however users can choose to use all edges if desired.

Whichever layout is used the final visualisation places the cluster nodes in a series of layers where each layer is a different clustering resolution and edges show the transition of samples through those resolutions. Edges are coloured according to the number of samples they represent and the in-proportion metric is used to control the edge transparency, highlighting more important edges. By default, the size of nodes is adjusted according to the number of samples in the cluster and their colour indicates the clustering resolution. The clustree package also includes options for controlling the aesthetics of nodes based on the attributes of samples in the clusters they represent as shown in the following examples.

While a clustering tree is conceptually similar to the tree produced through hierarchical clustering there are some important differences. The most obvious are that a hierarchical clustering tree is the result of a particular clustering algorithm and shows the relationships between individual samples while the clustering trees described here are independent of clustering method and show relationships between clusters. The branches of a hierarchical tree show how the clustering algorithm has merged samples. In contrast, edges in a clustering tree show how samples move between clusters as the resolution changes and nodes may have multiple parents. While it is possible to overlay information about samples on a hierarchical tree this is not commonly done but is a key feature of the clustree package and how clustering trees could be used in practice.

## A demonstration using simulations

To demonstrate what a clustering tree can look like in different situations and how it behaves as a dataset is over-clustered we present some illustrative examples using simple simulations (see methods). We present five scenarios: random uniform noise (Simulation A), a single cluster (Simulation B), two clusters (Simulation C), three clusters (Simulation D) and four clusters (Simulation E). Each cluster consists of 1000 samples (points) generated from a 100 dimensional normal distribution and each synthetic dataset has been clustered using *k-*means clustering with *k* = 1,*…*, 8. We then use the clustree package to produce clustering trees for each dataset (Figure 2).

**Figure 2:**
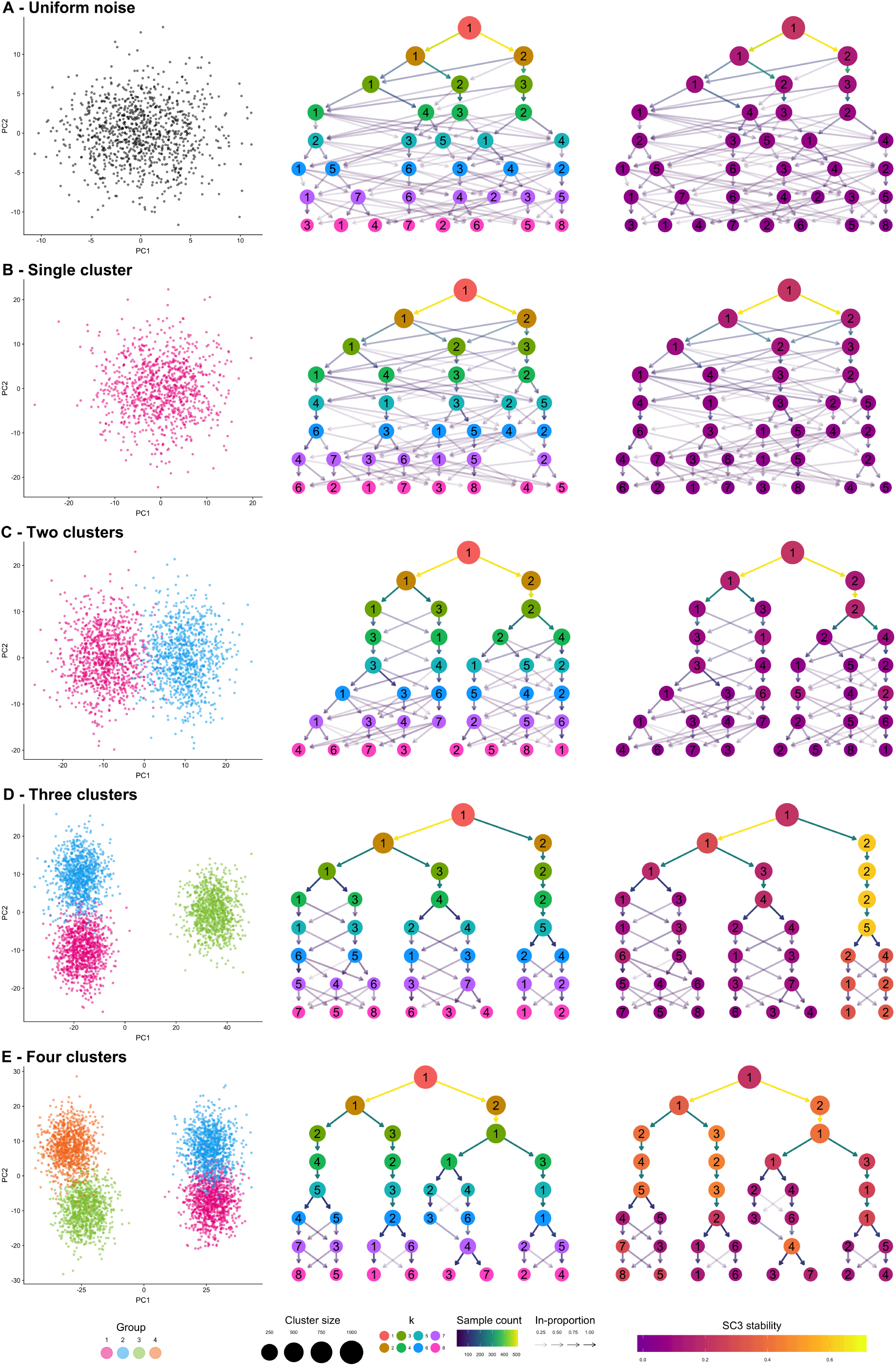
Five synthetic datasets used to demonstrate clustering trees. For each dataset a scatter plot of the first two principal components, a default clustering tree and and clustering tree with nodes coloured by the SC3 stability index from purple (lowest) to yellow (highest) are shown. The five datasets contain: A) random uniform noise, B) a single cluster, C) two clusters, D) three clusters and E) four clusters.

Looking at the first two examples (uniform noise (Figure 2A) and a single cluster (Figure 2B)) we can clearly see how a clustering tree behaves when a clustering algorithm returns more clusters than are truly present in a dataset. New clusters begin to form from multiple existing clusters and many samples switch between branches of the tree resulting in low in-proportion edges. Unstable clusters may also appear then disappear as the resolution increases as seen in Figure 2E. As we add more structure to the datasets the clustering trees begin to form clear branches and low in-proportion edges tend to be confined to sections of the tree. By looking at which clusters are stable and where low in-proportion edges arise we can infer which areas of the tree are likely to be the result of true clusters and which are caused by over-clustering.

The second clustering tree for each dataset shows nodes coloured according to the SC3 stability index for each cluster. As we would expect in the first two examples no cluster receives a high stability score. However, while we clearly see two branches in the clustering tree for the two cluster example (Simulation C) this is not reflected in the SC3 scores. No cluster receives a high stability score, most likely due to the high number of samples moving between clusters as the resolution increases. As there are more true clusters in the simulated datasets the SC3 stability scores become more predictive of the correct resolution to use, however it is important to look at the stability scores of all clusters at a particular resolution as taking the highest individual cluster stability score could lead to the incorrect resolution being used, as can be seen in the four cluster example (Simulation E). These examples show how clustering trees can be used to display existing clustering metrics in a way that can help to inform parameter choices.

## A simple example

To further illustrate how a clustering tree is built, we will work through an example using the classical iris dataset [16]. This dataset contains measurements of the sepal length, sepal width, petal length and petal width from 150 iris flowers, 50 from each of three species: *Iris setosa, Iris versicolor* and *Iris virginica*. The iris dataset is commonly used as example for both clustering and classification problems with the *Iris setosa* samples being significantly different to, and linearly separable from, the other samples. We have clustered this dataset using *k-*means clustering with *k =* 1*,…*, 5 and produced the clustering tree shown in Figure 3A.

**Figure 3:**
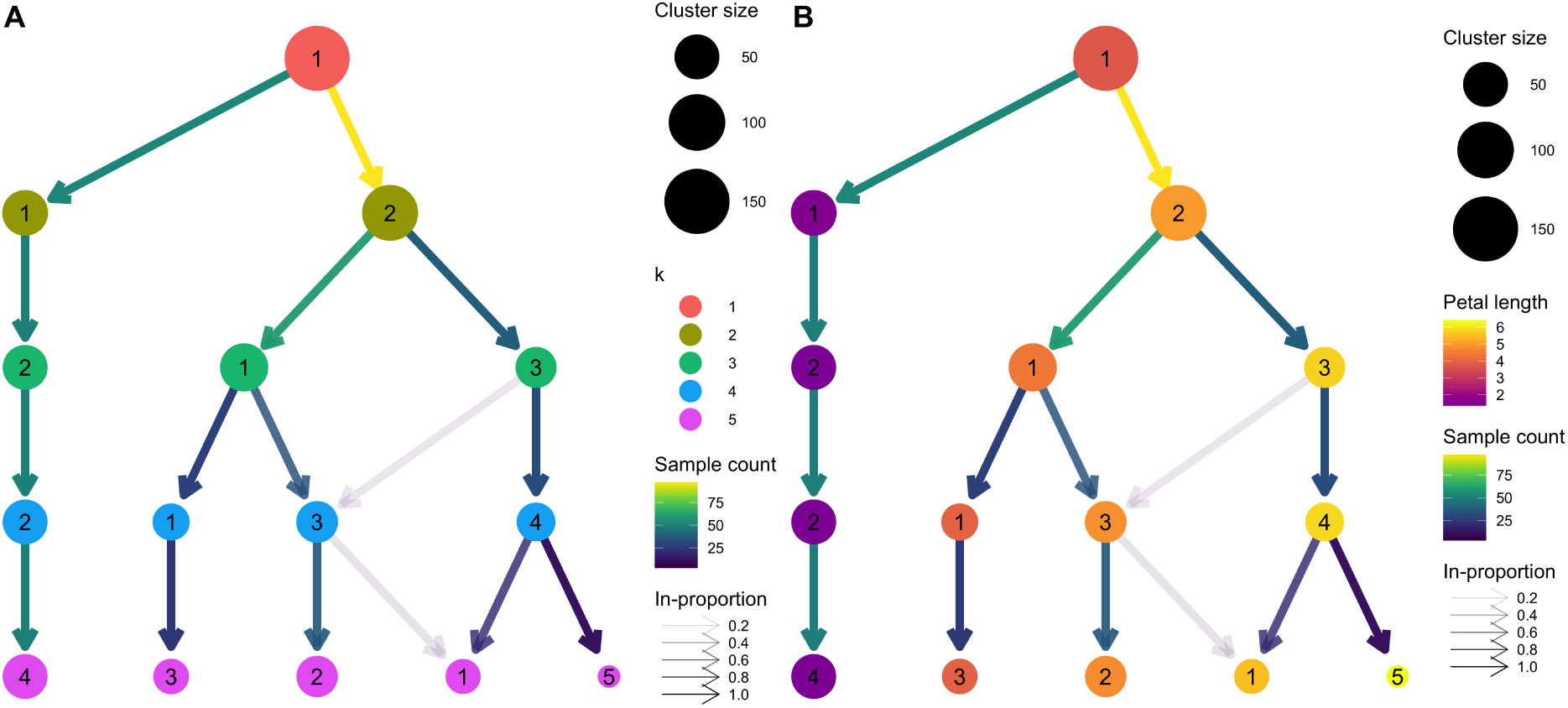
Clustering trees based on *k-*means clustering of the iris dataset. In A, nodes are coloured according to the value of *k* and sized according to the number of samples they represent. Edges are coloured according to the number of samples (from blue representing few to yellow representing many) and the transparency adjusted according to the in-proportion, with stronger lines showing edges that are more important to the higher resolution cluster. Cluster labels are randomly assigned by the *k-*means algorithm. B shows the same tree with the node colouring changed to show the mean petal length of the samples in each cluster.

We see that there is one branch of the tree that is clearly distinct (presumably representing *Iris setosa*), remaining unchanged regardless of the number of clusters. On the other side we see the cluster at *k* = 2 cleanly splits into two clusters (presumably *Iris versicolor* and *Iris virginica*) at *k* = 3 but as we move to *k* = 4 and *k* = 5 we see clusters being formed from multiple branches with more low in-proportion edges. As we have seen in the simulated examples, this kind of pattern can indicate that the data has become over-clustered and we have begun to introduce artificial groupings.

We can check our assumption that the distinct branch represents the *Iris setosa* samples and the other two clusters at *k* = 3 are *Iris versicolor* and *Iris virginica* by overlaying some known information about the samples. In Figure 3B we have coloured the nodes by the mean petal length of the samples they contain. We can now see that clusters in the distinct branch have the shortest petals, with Cluster 1 at *k* = 3 having an intermediate length and Cluster 3 the longest petals. This feature is known to separate the samples into the expected species with *Iris setosa* having the shortest petals on average, *Iris versicolor* an intermediate length and *Iris virginica* the longest.

Although this is a very simple example it highlights some of the benefits of viewing a clustering tree. We get some indication of the correct clustering resolution by examining the edges and we can overlay known information to assess the quality of the clustering. For example, if we observed that all clusters had the same mean petal length it would suggest that the clustering has not been successful as we know this is an important feature that separates the species. We could potentially learn more by looking at which samples follow low proportion edges or overlaying a series of features to try and understand what causes particular clusters to split.

## Clustering trees for single-cell RNA-seq data

One field that has begun to make heavy use of clustering techniques is the analysis of single-cell RNA-sequencing (scRNA-seq) data. Single-cell RNA-sequencing is a recently developed technology that can measure how genes are expressed in thousands to millions of individual cells [18]. This technology has been rapidly adopted in fields like developmental biology and immunology where it is valuable to have information from single cells rather than measurements that are averaged across the many different cells in a sample using older RNA sequencing technologies. One of the key uses for scRNA-seq is to discover and interrogate the different cell types present in a sample of a complex tissue. In this situation, clustering is typically used to group similar cells based on their gene expression profiles. Differences in gene expression between groups can then be used to infer the identity or function of those cells [19]. The number of cell types (clusters) in an scRNA-seq dataset can vary depending on factors such as the tissue being studied, its developmental or environmental state and the number of cells captured. Often the number of cells types is not known before the data is generated and some samples can contain dozens of clusters. Therefore, deciding which clustering resolution to use is an important consideration in this application.

As an example of how clustering trees can be used in the scRNA-seq context we consider a commonly used Peripheral Blood Mononuclear Cell (PBMC) dataset. This dataset was originally produced by 10x Genomics and contains 2700 peripheral blood monocuclear cells, representing a range of well-studied immune cell types [20]. We have analysed this dataset using the Seurat package [21], a commonly used toolkit for scRNA-seq analysis, following the instructions in their tutorial with the exception of varying the clustering resolution parameter from zero to five (see methods). Seurat uses a graph-based clustering algorithm and the resolution parameter controls the partitioning of this graph, with higher values resulting in more clusters. The clustering trees produced from this analysis are shown in Figure 4.

**Figure 4:**
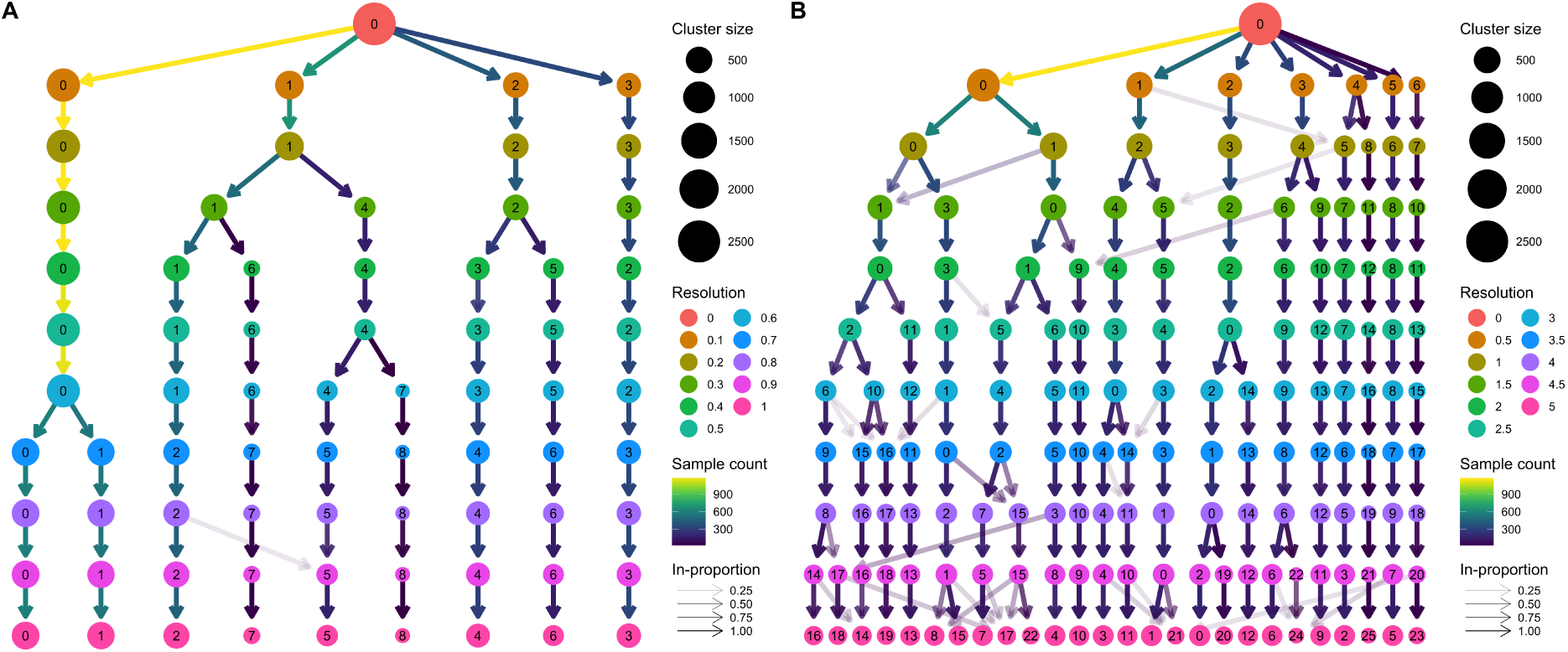
Two clustering trees of a dataset of 2700 Peripheral Blood Mononuclear Cells (PBMCs). A) results from clustering using Seurat with resolution parameters from zero to one. At a resolution of 0.1 we see the formation of four main branches, one of which continues to split up to a resolution of 0.5, after which there are only minor changes. B) resolutions from zero to five. At the highest resolutions we begin to see many low in-proportion edges indicating cluster instability. Seurat labels clusters according to their size with Cluster 0 being the largest.

The clustering tree covering resolutions zero to one in steps of 0.1 (Figure 4A) shows that four main branches form at a resolution of just 0.1. One of these branches, starting with Cluster 3 at resolution 0.1, remains unchanged while the branch starting with Cluster 2 splits only once at a resolution of 0.4. Most of the branching occurs in the branch starting with Cluster 1 which consistently has sub-branches split off to form new clusters as the resolution increases. There are two regions of stability in this tree; at resolution 0.5-0.6 and resolution 0.7-1.0 where the branch starting at Cluster 0 splits in two.

Figure 4B shows a clustering tree with a greater range of resolutions, from zero to five in steps of 0.5. By looking across this range we can see what happens when the algorithm is forced to produce more clusters than are likely to be truly present in this dataset. As over-clustering occurs we begin to see more low in-proportion edges and new clusters forming from multiple parent clusters. This suggests that those areas of the tree are unstable and that the new clusters being formed are unlikely to represent true groups in the dataset.

Known marker genes are commonly used to identify the cell types that specific clusters correspond to. Overlaying gene expression information onto a clustering tree provides an alternative view that can help to indicate when clusters containing pure cell populations are formed. Figure 5 shows the PBMC clustering tree in Figure 4A overlaid with the expression of some known marker genes.

**Figure 5:**
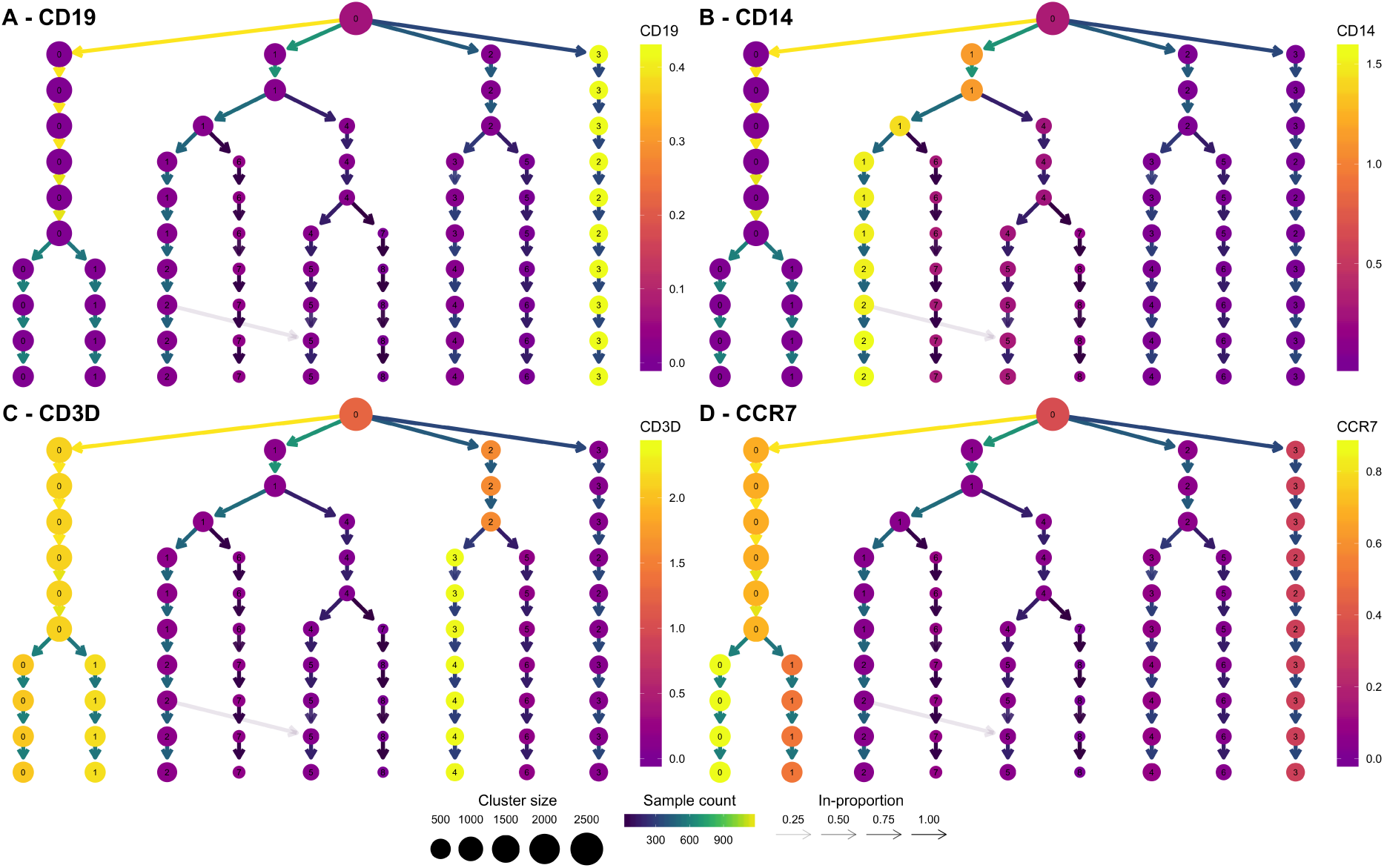
Clustering trees of the PBMC dataset coloured according to the expression of known markers. The node colours indicate the average of the log 2 gene counts of samples in each cluster. CD19 (A) identifies B cells, CD14 (B) shows a population of monocytes, CD3D (C) is a marker of T cells and CCR7 (D) shows the split between memory and naive CD4 T cells.

By adding this extra information, we can quickly identify some of the cell types. CD19 (Figure 5A) is a marker of B cells and is clearly expressed in the most distinct branch of the tree. CD14 (Figure 5B) is a marker of a type of monocyte, which becomes more expressed as we follow one of the central branches, allowing us to see which resolution identifies a pure population of these cells. CD3D (Figure 5C) is a general marker of T cells and is expressed in two separate branches, one which splits into low and high expression of CCR7 (Figure 5D), separating memory and naive CD4 T cells. By adding expression of known genes to a clustering tree, we can see if more populations can be identified as the clustering resolution is increased and if clusters are consistent with known biology. For most of the Seurat tutorial a resolution of 0.6 is used, but the authors note that by moving to a resolution of 0.8, a split can be achieved between memory and naive CD4 T cells. This is a split that could be anticipated by looking at the clustering tree with the addition of prior information.

## Discussion and conclusion

Clustering similar samples into groups is a useful technique in many fields, but often analysts are faced with the tricky problem of deciding which clustering resolution to use. Traditional approaches to this problem typically consider a single cluster or sample at a time and may rely on prior knowledge of sample labels. Here we present clustering trees, an alternative visualisation that shows the relationships between clusterings at multiple resolutions. While clustering trees cannot directly suggest which clustering resolution to use they can be a useful tool for helping to make that decision, particularly when combined with other metrics or domain knowledge.

Clustering trees display how clusters are divided as resolution increases, which clusters are clearly separate and distinct, which are related to each other and how samples change groups as more clusters are produced. Although clustering trees can appear similar to the trees produced from hierarchical clustering there are several important differences. Hierarchical clustering considers the relationships between individual samples and doesn’t provide an obvious way to form groups. In contrast, clustering trees are independent of any particular clustering method and show the relationships between clusters, rather than samples, at different resolutions, any of which could be used for further analysis.

To illustrate the uses of clustering trees we presented a series of simulations and two examples of real analyses, one using the classical iris dataset and a second based on a complex scRNA-seq dataset. Both examples demonstrate how a clustering tree can help inform the decision of which resolution to use and how overlaying extra information can help to validate those clusters. This is of particular use to scRNA-seq analysis as these datasets are often large, noisy and contain an unknown number of cell types or clusters.

Even when determining the number of clusters is not a problem, clustering trees can be a valuable tool. They provide a compact, information dense, visualisation that can display summarised information across a range of clusters. By modifying the appearance of cluster nodes based on attributes of the samples they represent, clusterings can be evaluated and identities of clusters established. Clustering trees potentially have applications in many fields and in the future could be adapted to be more flexible, such as by accommodating fuzzy clusterings. There may also be uses for more general clustering graphs to combine results from multiple sets of parameters or clustering methods.

## Methods

### clustree

The clustree software package is built for the R statistical programming language. It relies on the ggraph package (https://github.com/thomasp85/ggraph), which is itself built on the ggplot2 [22] and tidygraph packages (https://github.com/thomasp85/tidygraph). Clustering trees are displayed using the Reingold-Tilford tree layout or the Sugiyama layout, both available as part of the igraph package.

### Simulations

Simulated datasets were constructed by generating points from statistical distributions. The first simulation (Simulation A) consists of 1000 points randomly generated from a 100 dimensional space using a uniform distribution between zero and 10. Simulation B consists of a single normally distributed cluster of 1000 points in 100 dimensions. The centre of this cluster was chosen from a normal distribution with mean zero and standard deviation 10. Points were then generated around this centre from a normal distribution with mean equal to the centre point and a standard deviation of five. The remaining three simulations were produced by adding additional clusters. In order to have a known relationship between clusters the centre for the new clusters was created by manipulating the centres of existing clusters. For Cluster 2 a random 100 dimensional vector was generated from a normal distribution with mean zero and standard deviation two and added to the centre for Cluster 1. Centre 3 was the average of Centre 1 and Centre 2 plus a random vector from a normal distribution with mean zero and standard deviation five. To ensure a similar relationship between clusters 3 and 4 as between clusters 1 and 2, Centre 4 was produced by adding half the vector used to produce Centre 2 to Centre 3 plus another vector from a normal distribution with mean zero and standard deviation two. Points for each cluster were generated in the same way as for Cluster 1. Simulation C consists of the points in clusters 1 and 2, Simulation D consists of clusters 1, 2 and 3, Simulation E consists of clusters 1, 2, 3 and 4. Each simulated dataset was clustered using the “kmeans” function in the stats package with values of *k* from one to eight, a maximum of 100 iterations and 10 random starting positions. The clustering tree visualisations were produced using the clustree package with the tree layout. The simulated datasets and the code use to produce them are available from the repository for this paper (https://github.com/Oshlack/clustree-paper).

### Iris dataset

The iris dataset is available as part of R. We clustered this dataset using the “kmeans” function in the stats package with values of *k* from one to five. Each value of *k* was clustered with a maximum of 100 iterations and with 10 random starting positions. The clustree package was used to visualise the results using the Sugiyama layout. The clustered iris dataset is available as part of the clustree package.

### PBMC dataset

The PBMC dataset was downloaded from the Seurat tutorial page (http://satijalab.org/seurat/pbmc3k_tutorial.html) and this tutorial was followed for most of the analysis. Briefly cells were filtered based on the number of genes they express and the percentage of counts assigned to mitochondrial genes. The data was then log-normalised and 1838 variable genes identified. Potential confounding variables (number of unique molecular identifiers and percentage mitochondrial expression) were regressed from the dataset before performing principal component analysis on the identified variable genes. The first 10 principal components were then used to build a graph which was partitioned into clusters using Louvain modularity optimisation [23] with resolution parameters in the range zero to five, in steps of 0.1 between zero and one and then in steps of 0.5. Clustree was then used to visualise the results using the tree layout.

## Declarations

### Ethics

Not applicable.

### Availability of data and materials

The clustree package (RRID: SCR_016293) is available from CRAN (https://CRAN.R-project.org/package=clustree) and is being developed on GitHub at https://github.com/lazappi/clustree. The code and datasets used for the analysis in this paper are available from https://github.com/Oshlack/clustree-paper. The clustered iris dataset is included as part of clustree and the PBMC dataset can be downloaded from the Seurat tutorial page (http://satijalab.org/seurat/pbmc3k_tutorial.html) or the paper GitHub repository.

## Acknowledgements

Thank you to Marek Cmero for providing comments on a draft of the manuscript.

